# Three simple steps to improve the interpretability of EEG-SVM studies

**DOI:** 10.1101/2021.12.14.472588

**Authors:** Coralie Joucla, Damien Gabriel, Juan-Pablo Ortega, Emmanuel Haffen

## Abstract

Research in machine-learning classification of electroencephalography (EEG) data offers important perspectives for the diagnosis and prognosis of a wide variety of neurological and psychiatric conditions, but the clinical adoption of such systems remains low. We propose here that much of the difficulties translating EEG-machine learning research to the clinic result from consistent inaccuracies in their technical reporting, which severely impair the interpretability of their often-high claims of performance. Taking example from a major class of machine-learning algorithms used in EEG research, the support-vector machine (SVM), we highlight three important aspects of model development (normalization, hyperparameter optimization and cross-validation) and show that, while these 3 aspects can make or break the performance of the system, they are left entirely undocumented in a shockingly vast majority of the research literature. Providing a more systematic description of these aspects of model development constitute three simple steps to improve the interpretability of EEG-SVM research and, in fine, its clinical adoption.

## Introduction

Electroencephalography (EEG), and its associated methodologies such as evoked/event-related potentials (EP/ERP), allows recording the electric fields generated by neuronal activity non-invasively simply by placing electrodes on the scalp. This simplicity makes EEG possibly the most widespread tool in human brain research, with significant impact in all fields of psychology and cognitive neuroscience. However, while its recording is relatively simple, identifying valuable information within the various waveforms of the EEG remains a complex technical process, as these signals are noisy, complex and of high dimensionality (Cohen, 2017). In effect, this analytical complexity limits the routine indication of EEG/ERP in the clinic to, besides cerebral death exams, only a handful of conditions incl. epilepsy, sleep disorders and disorders of consciousness (Quigg, Shneker & Domer, 2001).

In an attempt to increase the translatability between EEG/ERP research and the clinic, recent years have seen a surge in the development of machine-learning systems aiming to standardize and automate the classification of EEG signals into mental/physiological or pathological states of interest. These systems offer the promise to move from group-level statistics to patient-by-patient decidability, not only in traditional indications such as sleep (Fiorillo et al., 2019), consciousness (Engemann et al., 2018) or epilepsy (Siddiqui et al., 2020), but also in a wider number of major central nervous system disorders including Alzheimer’s disease, schizophrenia, depression, and pain (Leiser et al. 2011).

Yet, despite years of research, the clinical adoption of AI-related technologies in EEG medical diagnostics remain slow (Kelly et al. 2019) and is in stark contrast with the high-to-very-high performance indicators (e.g. diagnostics in the high-90% accuracies) seemingly promised for such systems by the specialized research literature. While reasons for this slow adoption may include human factor and regulatory aspects (Elkefi, Wang & Asan, 2020), we propose here that much of the difficulties translating EEG-machine learning research to the clinic in fact result from consistent inaccuracies in their technical reporting, which severely impair the interpretability of their often-high claims of performance. Taking example from a major class of machine-learning algorithms used in EEG research, the support-vector machine (SVM; Cortes & Vapnik, 1995), we review three important degrees-of-freedom that researchers have in developing and evaluating such algorithms (normalization, hyperparameter optimization and cross-validation) and show that, while these 3 aspects can make or break the performance of the system, they are left entirely undocumented in a shockingly vast majority of the research literature. In our view, providing a more systematic description of these aspects of model development constitute three simple steps to improve the interpretability of EEG-SVM research and, in fine, its clinical adoption.

### A suspiciously positive distribution of accuracy in the EEG-SVM literature

We queried the pubmed database on Jan. 14th 2021 for articles covering SVM classification of EEG signals, excluding articles focussing on detecting EEG microstructural elements in sleep (e.g. spindles) and epilepsy (e.g. spikes). The search equation was the following : “eeg support vector machine classification NOT epilepsy NOT seizure NOT review NOT sleep”. The query returned an initial list of 434 articles, to which 4 articles were added from additional sources. The eligibility of the selected papers was established by first examining the titles, then the abstracts, and finally the entire articles. 262 articles that did not satisfy the criteria were rejected, leaving 176 articles included for the present analysis.

Publication dates revealed that this field of research is recent and active, starting in the early 2010’s and peaking after 2015; in 2018 alone, the number of publications was equivalent to the number of articles published in 2015 and 2016 combined (Figure 1B). Geographically (Figure 1A), the majority of studies came from China (n=51, 28.98% of first author affiliations), USA (n=9, 5.11%) and India (n=8, 4.55%) (note: that distribution can be compared with a similar map of EEG deep-learning research - Roy et al. 2019), and focused on classifying emotional and cognitive brain states (75 articles); the remaining 55 and 36 articles concerned brain-computer interface (BCI) and clinical diagnosis respectively. For each included article, 36 research items were then extracted describing authorship and publication metrics, type of input data, EEG pre-processing, SVM methodology, results obtained and availability of code and data. In the following, we focus on items related to SVM methodology and how they relate to authorship, model performance and reproducibility; we nevertheless provide the full data table in Supplementary material.

**Figure 1.**
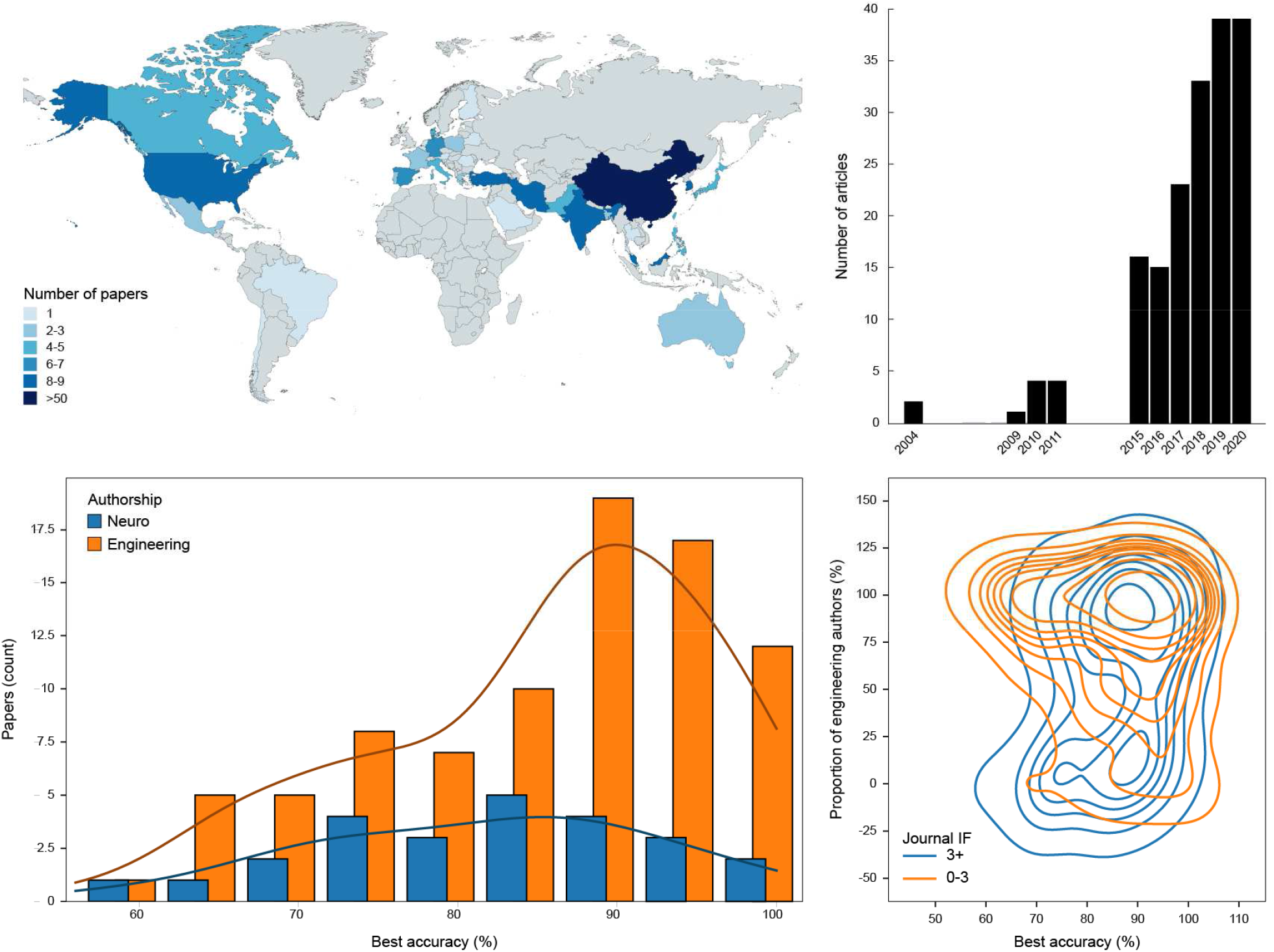
The recent EEG-SVM research literature reports distributions of percentage accuracy that are suspiciously positive for such a variety of research problems (n=176). **(A)** Map of country of first author affiliation. **(B)** Yearly counts of articles included in our review. **(C)** Articles published in below-median IFs (orange) are predominantly published by authors affiliated with computer science or engineering departments, and report distributions of accuracies that are more positively skewed and more peaky in the 90-100% range. **(D)** Comparison of the accuracies reported by articles solely published by authors affiliated with natural science departments (blue; n=25) and articles published solely by authors affiliated with engineering (orange; n=90)

157 of the 176 articles reported performance metrics in the form of two-class classification accuracy. With a median of 87.8%, the distribution of accuracy values across papers was strongly skewed towards high-to-very-high performance (Figure 1C). Articles solely published by authors affiliated with natural science departments (N=25, 15%) had a more uniform distribution of reported accuracy than articles with authors affiliated with computer science or engineering departments (N=90, 61%), which feature increasingly peaky distributions in the high-80-90% as the proportion of these authors increase (Figure 1C). This disciplinary effect is partly confounded with journal impact factors (median IF=3.1), with articles published in below-median IFs being predominantly authored by CS/Engineering authors and reporting higher accuracies (Mdn=88.89%) than articles in above-median IFs (Mdn=85.96%; Figure 1D).

Such distributions of high accuracies are unlikely to reflect the true distribution of performance over the population of all EEG-SVM classification problems, especially across domains as varied as emotion classification, brain-computer interfaces or psychiatric diagnosis. While these distributions likely incorporate biases in topic selection (in which researchers tend to avoid automatic classification problems for which the link between EEG signals and the cognitive/clinical variables of interest is expected to be poor) and in result publication (in which authors and journals only publish studies that “work” - Chavalarias et al., 2016), the fact that their positive skew increases with the technicity of authors (as measured by departmental affiliation) also suggests that the technicalities of how to build and evaluate EEG-SVM systems constitute “researchers’ degrees of freedom” (Simmons, Nelson & Simonsohn, 2011) that, in fact, allow optimizing results to almost-arbitrary levels of performance.

However, in our view, this does not suggest that a majority of these studies are flawed or misrepresenting the strength of the association of EEG to symptoms. What this suggests, rather, is that the performance of a given EEG-SVM classification system is impossible to interpret without complete disclosure and understanding of how that system was built and evaluated. In the following, we present three important aspects of system building and evaluation (normalization, hyperparameter optimization and cross-validation) which strike us as particularly important, both because they critically affect the interpretation of the reported performance and because they are undocumented in a majority of the literature, and give recommendations on how these items should be reported in research articles.

## Three simple steps to improve the interpretability of EEG-SVM studies

### Normalization

SVM classification works by mapping training examples, represented as multidimensional data points in a vector space (Figure 2), into a higher-dimensional space where the decision problem between the classes is easier (linearly-separable). This mapping is based on weighting functions, or kernels, which typically assume that all the data being compared are within a standard range. It is therefore very important to normalise the feature vectors before training the classifier.

**Figure 2.**
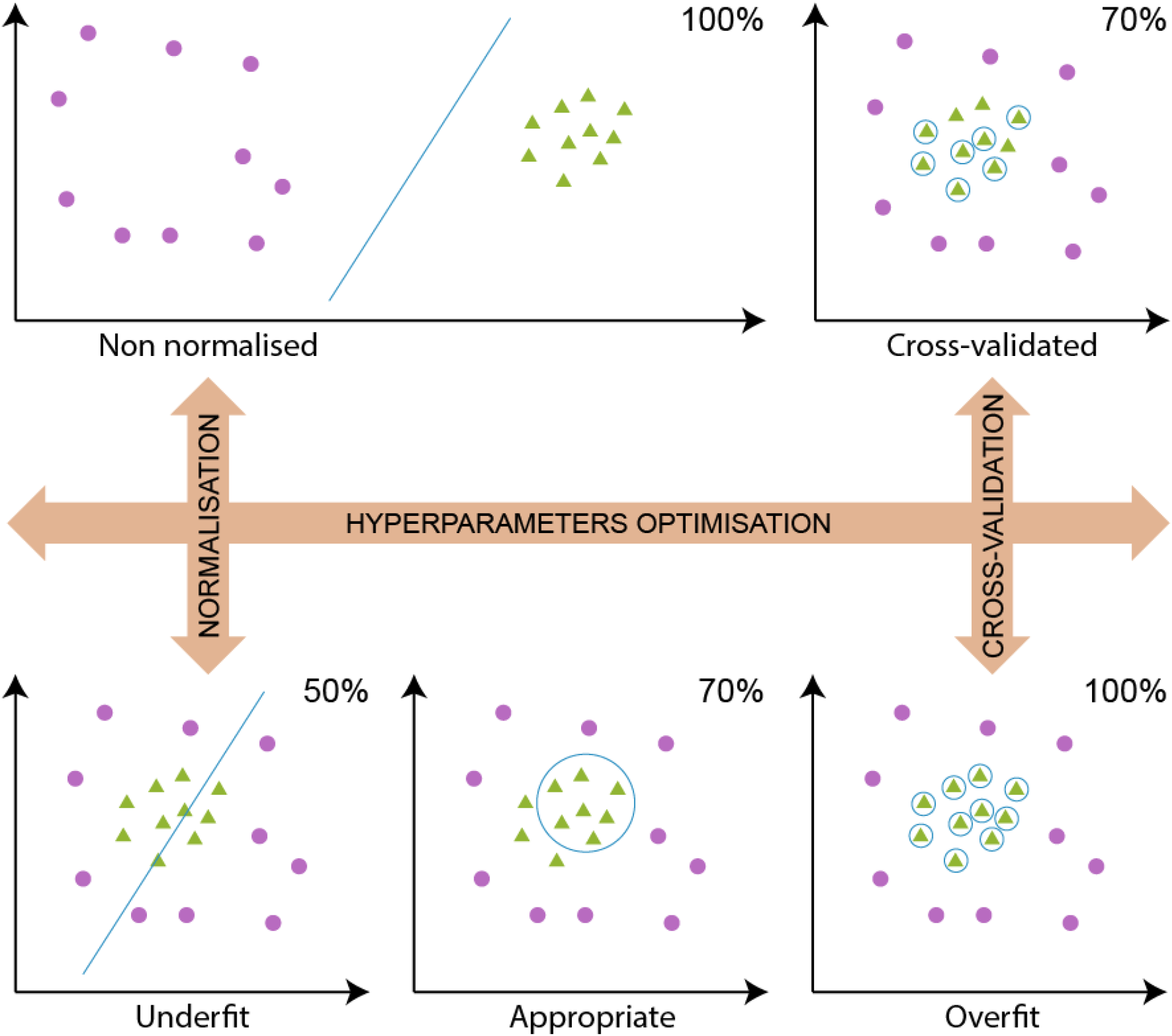
Three important researcher’s degrees of freedom in building and evaluating EEG-SVM classification systems, which can make or break system performance. Left, vertically: Normalisation. If data have different scales (e.g. patient data with larger EEG impedances than controls), failing to normalize can make a difficult problem (bottom) appear spuriously easy (top). **Horizontally**: Hyperparameter optimization. Model parameters control, for instance, the tradeoff between the accurate classification of the training examples (improving from left to right) and the ability to generalize to new, unseen examples (improving from right to left). **Right, vertically:** Cross-validation. Testing on the same dataset used for training can make an overfitted model appear perfectly accurate (bottom), while testing on new, unseen data reveals that its generalization ability is in fact poor (top).

Normalization issues can make or break SVM classification results. For instance, if multiple dimensions in the data have very different ranges (e.g. different power levels due to electrode position; or voltage data in *μ*volts next to spectrum power data in dBs), failing to normalize data within each dimension will result in the model overemphasizing the numerical contribution of one dimension compared to the other, and possibly degrade performance. Conversely, if data from one’s classes of interest have different scales or offsets (e.g. patient data with larger EEG impedances than controls), failing to normalize data in all dimensions can make a difficult problem appear spuriously linearly-separable (Figure 2-left)

Despite its importance, we found that the proportion of articles for which no normalization is mentioned is overwhelming: 71.0% (Figure 3-top). Different normalization methods exist: one can e.g. match the minimum and maximum values of each dimension to 0 and 1, or to -1 and 1 (min-max normalisation), or subtract the mean of the values and divide by the standard deviation (z-score). Of the remaining articles, 13.6% opted for z-score normalisation, and the rest are spread over variants of min-max normalisation.

**Figure 3.**
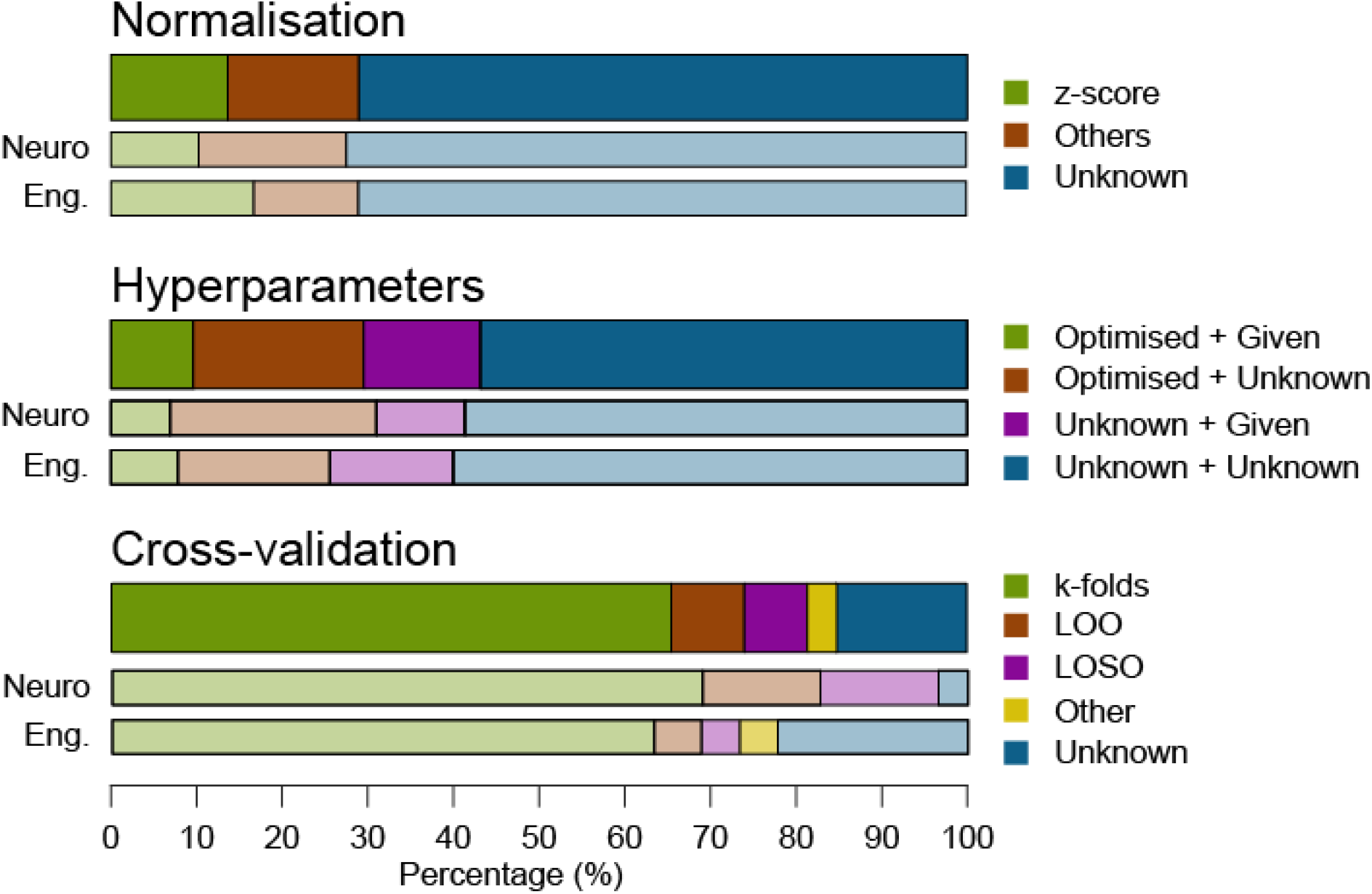
Three critical aspects of EEG-SVM system building and evaluation are left undocumented in a majority of the research literature. (Top: normalization; middle: hyperparameter optimization; bottom: cross-validation). These proportions did not depend on the disciplinary background of the study authors

An accurate description of the process of normalizing data should not only describe its mathematical method, but also explain over what set of data it is done: for instance, normalizing data separately within patients and controls would solve the problem illustrated in Figure 2-left, but normalizing across all data wouldn’t. The following example, adapted from one of our reviewed articles (Zarei et al.2017) illustrates what a transparent, useful description of normalization can look like:

> *“2*.*2*.*1. Normalization*
>
> **“***In this step, all EEG signals are preprocessed to have unit variance and zero mean to eliminate differences in power levels between the signals due to electrode position. The normalization process facilitates the PCA calculations to ensure equal weighting*. **[mention normalization, and explain the rationale for doing so in this specific problem]**
>
> *This process is done by subtracting the overall average from each EEG signal and dividing it by its standard deviation (Equ*.*1)*. **[give normalization method; formalize it with equation]**
>
> *As mentioned before, two datasets IVa and IVb from BCI Competition III are used in this study. Both dataset consist in EEG signals involving a MI task with two classes. [*…*] Each dataset for a subject is divided into 50/50 fractions for training and testing sets for evaluation of our proposed methods. Therefore, each training set and testing set for a subject contains*
>
> *59 channel data in each class. All channel data in both training and testing sets are normalized separately using Equ*.*1 (see Fig.1 - Block diagram)”*. **[explain over what set of data the normalization is applied; illustrate with block diagram]**

### Hyperparameter optimization

SVMs predict the belonging of a trial (e.g., a patient’s epoch of EEG recording in response to a stimulus) to a given class of data (e.g., pathological/not) by constructing a model of the boundary between classes. To do so, SVMs use a weighting function, or kernel, to map training examples (trials, and their annotated class) to points in a space constructed to maximise the width of the gap between the two classes (Cortes & Vapnik, 1995). New examples are then mapped into that same space and predicted to belong to one or the other class depending on which side of the gap they fall into.

A variety of kernels are used to construct SVMs, such as linear and radial basis function (RBF) kernels, and their behaviour can typically be adjusted using hyperparameters that regulate what the model optimizes for (Herbrich, 2001). For instance, parameter *C* controls the tradeoff between the accurate classification of training examples and the maximization of the decision-boundary margin, i.e. how much one is willing to compromise on exactitude for the benefit of generalizability (Figure 2-bottom).

The choice of kernel hyperparameters strongly influences the performance of the model and it is highly recommended to optimize them (e.g. by testing for all possible values in a systematic “grid search”) for one’s specific classification problem. Yet, out of the 176 articles included in this study, 124 (70.5%) did not mention any hyperparameter optimization, and 135 (76.7%) did not provide the selected hyperparameters (Figure 3-middle). Only 76 studies (43.2%) optimised and/or provided the coefficients of the kernels used (59.6% of studies who explicitly searched for optimal parameters used grid search). This indicates that either the authors were not aware of the importance of these elements and did not employ them, or that they used a particular method or parameters but did not see fit to document it.

An accurate description of the process of optimizing hyperparameters should explain what kernel is used, what hyperparameters are optimized (e.g. C), with what method (e.g. grid search) and which value is retained. As for normalization, it should also explain on what set of data these parameters are optimized: the normal practice is to optimize parameters on a different set of data than the one used for evaluation, and test whether these values generalize well to unseen data (a process made even more complicated by cross-validation, see below). The following example, adapted from Arvaneh et al. (2011), Hou et al. (2020) and Kirar & Agrawal (2018), illustrates what a transparent, useful description of hyperparameter optimization can look like:

> *“The Radial Basis Function (RBF) was used as the kernel function of the SVM, as suggested in [30] for problems in which, like here, the number of instances far exceeds the number of features*. ***[say what kernel is used, and why]***
>
> *To achieve best performance, kernel hyperparameters C (regularization) and g (radius) were optimized by grid search, using 10-fold cross-validation on the training data*. ***[say what hyperparameters are optimized and on what dataset]***
>
> *The search domains were [1-500] for C and [1-100] for g, with a step size of 10”*. ***[give details of optimization method]*** *“Following optimization, the values retained for testing were C = 100 and g = 10*.*”* ***[give hyperparameter values]***

### Cross-validation

When translating a machine-learning system to clinical practice, the only performance metrics that really matters is its ability to generalize to new, unseen examples. To evaluate this ability, it is common to divide the available data into a set for training the model and a set for testing it. One can do this only once or, for better statistical estimation of the underlying sampling distribution, split data into different partitions that are then used successively as training or testing sets - a process called cross-validation (Wong, 2015). The most popular procedure is called k-fold cross-validation, in reference to the number of partitions that data set is to be split into. Leave-One-Out (LOO) is an alternative method where, for each sample in the dataset, a model is built using all other samples and then tested on the selected sample. Leave-One-Subject-Out (LOSO) is the same procedure as LOO, but where data from a whole participant is set aside (for a comparison of these methods, see e.g. Simon 2007).

Not using cross-validation (i.e. testing on training data) systematic over-evaluates model performance, in particular in cases where model parameters are adjusted to overfit training data (Figure 2-right) and where accuracy can reach 100% (as reported, incidentally, in n=6 articles…). Even when using a cross-validation procedure, system generalizability can be overestimated when model hyperparameters are optimized based on testing data (i.e. training with successive values of the parameter, and selecting both the optimal value and the final system accuracy by cross-validation on the same data). The normal practice in this case should be to further split training data into “training” and “validation” sets, the latter being used to select the best hyperparameter, and to leave it to the “test” set to measure the performance of the best model. Park & Han (2018) note in passing that this usage of the word ‘validation’ in machine-learning, which refers to the fine-tuning stage of model development, can create confusions in the fields of medicine and health, where it can be wrongly taken to mean verifying model performance (what the field instead called ‘testing’).

Worryingly, a sizeable n=27 (15.3%) of articles did not mention using cross-validation when evaluating model performance (Figure 3-bottom). When they did (n=149), a majority used k-fold cross-validation (65.3%; 10-fold: 34.7%), LOO (8.5%) or LOSO (7.4%). However, only n=28 articles (18.8%) of those reporting cross-validated performance mentioned using a separate validation set for model optimization

The following example, adapted from Jessen, Obleser & Tune (2021), illustrates what a transparent, useful description of cross-validation can look like:

> *“As a first step, we split the continuous data into two data segments, with 80% of the data reserved for training, and the remaining 20% set aside for final model testing*. ***[clearly describe the stratification of your data]***
>
> *To optimize hyperparameter λ, we then apply a technique called k-fold cross-validation. To this end, we split the training data into 4 equal sized segments, referred to as folds. Within the cross-validation routine, training and validation sets are rotated until each fold has served as validation set while the remaining three folds are jointly used for training (see Fig - schematic of data sets). In the next step, we average model performance (i*.*e*., *Pearson’s r and MSE) per tested λ value across folds to identify the λ value that yields the best model performance*. ***[say whether you use cross-validation, what method you use, and if you use a validation set to optimize model parameters*.]**
>
> *Finally, we apply this optimal λ parameter for model estimation using all training data and test it on the initially left-out test data segment. All results reported in the following correspond to performance estimated on such test data”*. ***[clearly state whether reported performance is computed on training, validation or testing data]***

## Conclusion

In sum, because they can utterly modify the meaning of a system’s reported accuracy, documenting the three aspects of normalization, hyperparameter optimization and cross-validation constitute simple and easy steps to improve the interpretability and clinical translatability of EEG-SVM research. Interestingly, while a majority of the research literature fails to do so, these proportions do not appear, at least for normalization and parameter optimization, to much depend on the disciplinary background of the study authors (Figure 3; note though that undocumented cross-validation is more common in articles solely published by engineering authors). This lack of transparency is all the more so striking as, typically, the same articles give a thorough description of the EEG methodology (recording, processing, etc.) used to generate the data (see e.g. Aydin et al., 2015; Bamatraf et al., 2016; Torabi et al., 2017).

Such recommendations of transparency overlap with a general call for greater science openness, not only for the access to publications but also to data and code. On that aspect too, our set of articles appears to be lacking: only 26.1% of our reviewed papers lists data as public or available on request, a proportion similar to medical journals like BMJ (Rowhani-Farid & Barnett, 2016) but smaller than journals with an explicit data-sharing policy, such as PLOS One (Federer et al. 2018) or, in the same technical field, than the 53% reported in a recent review of EEG & deep-learning research (Roy et al. 2019). Similarly, code sharing concerned a meager 3% of the articles, again a smaller proportion than what is now typical of fields with greater awareness of replicability such as psychological science and statistics (33%; Kucharský, Houtkoop & Visser, 2020). Initiatives to publish datasets and code with digital object identifier (DOI) such as the Open Science Framework (www.osf.io), which allow researchers to obtain credit for their data and to cite them, or journal submission procedure which explicitly incorporate data and code sharing, such as in the French Statistical Society’s Computo journal (https://computo.sfds.asso.fr), are possible levers to increase these trends.

Finally, the three aspects of model building highlighted here, while described in relation to the specific method of SVMs, are also examples of general concepts (feature selection, regularization, cross-validation) that are relevant for the greater corpus of EEG machine-learning research. For instance, a recent review of EEG deep-learning research (Roy et al. 2019) showed that an even greater proportion of articles did not use or mention hyperparameter search (80%) or cross-validation (42%. This suggests that, without the establishment of best practices for reporting such as advocated here and elsewhere (Poldrack, Huckins & Varoquaux, 2020), newer and increasingly sophisticated AI methods may even further degrade, rather than improve, the clinical translatability of the associated research results.

## Acknowledgements

Work funded by ANR SEPIA (2020-2024, to C.J). C.J. thanks JJ. Aucouturier (FEMTO-ST, CNRS / Université de Bourgogne Franche-Comté) for help finalizing the manuscript. The idea to include examples of possible paragraphs is inspired by Ekkekakis (2013).

## References

Arvaneh, M., Guan, C., Ang, K. K., & Quek, C. (2011). Optimizing the Channel Selection and Classification Accuracy in EEG-Based BCI. IEEE Transactions on Biomedical Engineering, 58(6), 1865–1873.

Aydin, S., Arica, N., Ergul, E., & Tan, O. (2015). Classification of Obsessive Compulsive Disorder by EEG Complexity and Hemispheric Dependency Measurements. International Journal of Neural Systems, 25(03), 1550010.

Bamatraf, S., Hussain, M., Aboalsamh, H., Qazi, E.-U.-H., Malik, A. S., Amin, H. U., Mathkour, H., Muhammad, G., & Imran, H. M. (2016). A System for True and False Memory Prediction Based on 2D and 3D Educational Contents and EEG Brain Signals. Computational Intelligence and Neuroscience, 2016, 1–11.

Chavalarias, D., Wallach, J. D., Li, A. H. T., & Ioannidis, J. P. (2016). Evolution of reporting P values in the biomedical literature, 1990-2015. Jama, 315(11), 1141–1148.

Cohen, M. X. (2017). Where does EEG come from and what does it mean?. Trends in neurosciences, 40(4), 208–218.

Cortes, C. & Vapnik, V. (1995). Support-vector networks. Machine learning, 20(3), 273–297.

Ekkekakis, P. (2013). The measurement of affect, mood, and emotion: A guide for health-behavioral research. Cambridge University Press.

Elkefi, S., Wang, H., & Asan, O. (2020). Organizational considerations from HFE to speed up the adoption of AI-related technology in medical diagnostics. In Proc. International Symposium on Human Factors and Ergonomics in Health Care, Vol. 9(1), pp. 230–234. Los Angeles, CA: SAGE Publications.

Engemann, D. A., Raimondo, F., King, J.-R., Rohaut, B., Louppe, G., Faugeras, F., Annen, J., Cassol, H., Gosseries, O., Fernandez-Slezak, D., Laureys, S., Naccache, L., Dehaene, S., & Sitt, J. D. (2018). Robust EEG-based cross-site and cross-protocol classification of states of consciousness. Brain, 141(11), 3179–3192.

Federer, L. M., Belter, C. W., Joubert, D. J., Livinski, A., Lu, Y. L., Snyders, L. N., & Thompson, H. (2018). Data sharing in PLOS ONE: an analysis of data availability statements. PloS one, 13(5), e0194768.

Fiorillo, L., Puiatti, A., Papandrea, M., Ratti, P.-L., Favaro, P., Roth, C., Bargiotas, P., Bassetti, C. L., & Faraci, F. D. (2019). Automated sleep scoring: A review of the latest approaches. Sleep Medicine Reviews, 48, 101204.

Herbrich, R. (2001). Learning kernel classifiers: Theory and algorithms. MIT Press.)

Hou, H.-R., Zhang, X.-N., & Meng, Q.-H. (2020). Odor-induced emotion recognition based on average frequency band division of EEG signals. Journal of Neuroscience Methods, 334, 108599.

Jessen, S., Obleser, J., & Tune, S. (2021). Neural Tracking in Infants–an Analytical Tool for Multisensory Social Processing in Development.

Kelly, C. J., Karthikesalingam, A., Suleyman, M., Corrado, G., & King, D. (2019). Key challenges for delivering clinical impact with artificial intelligence. BMC medicine, 17(1), 1–9.

Kirar, J. S., & Agrawal, R. K. (2018). Relevant Feature Selection from a Combination of Spectral-Temporal and Spatial Features for Classification of Motor Imagery EEG. Journal of Medical Systems, 42(5), 78.

Kucharský, Š., Houtkoop, B. L., & Visser, I. (2020). Code Sharing in Psychological Methods and Statistics: An Overview and Associations with Conventional and Alternative Research Metrics. OSF Preprints. February 24. doi:10.31219/osf.io/daews.

Leiser, S. C., Dunlop, J., Bowlby, M. R., & Devilbiss, D. M. (2011). Aligning strategies for using EEG as a surrogate biomarker: a review of preclinical and clinical research. Biochemical pharmacology, 81(12), 1408–1421.

Park, S. H., & Han, K. (2018). Methodologic guide for evaluating clinical performance and effect of artificial intelligence technology for medical diagnosis and prediction. Radiology, 286(3), 800–809.

Poldrack, R. A., Huckins, G., & Varoquaux, G. (2020). Establishment of best practices for evidence for prediction: a review. JAMA psychiatry, 77(5), 534–540.

Quigg, M., Shneker, B., & Domer, P. (2001). Current practice in administration and clinical criteria of emergent EEG. Journal of Clinical Neurophysiology, 18(2), 162–165.

Rowhani-Farid A, Barnett AG. Has open data arrived at the British Medical Journal (BMJ)? An observational study. BMJ Open. 2016;6: 1–8. pmid:27737882

Roy, Y., Banville, H., Albuquerque, I., Gramfort, A., Falk, T. H., & Faubert, J. (2019). Deep learning-based electroencephalography analysis: a systematic review. Journal of Neural Engineering, 16(5), 051001.

Siddiqui, M. K., Morales-Menendez, R., Huang, X., & Hussain, N. (2020). A review of epileptic seizure detection using machine learning classifiers. Brain Informatics, 7(1), 5.

Simmons, J. P., Nelson, L. D., & Simonsohn, U. (2011). False-positive psychology: Undisclosed flexibility in data collection and analysis allows presenting anything as significant. Psychological Science, 22(11), 1359–1366.

Simon, R. (2007). Resampling strategies for model assessment and selection. In Fundamentals of data mining in genomics and proteomics (pp. 173–186). Springer, Boston, MA.

Torabi, A., Daliri, M. R., & Sabzposhan, S. H. (2017). Diagnosis of multiple sclerosis from EEG signals using nonlinear methods. Australasian Physical & Engineering Sciences in Medicine, 40(4), 785–797.

Wong, T. T. (2015). Performance evaluation of classification algorithms by k-fold and leave-one-out cross validation. Pattern Recognition, 48(9), 2839–2846.

Zarei, R., He, J., Siuly, S., & Zhang, Y. (2017). A PCA aided cross-covariance scheme for discriminative feature extraction from EEG signals. Computer methods and programs in biomedicine, 146, 47–57.

